# Ascending and Descending Aortic ECM Hydrogels for Modeling Aortic Wall Biology

**DOI:** 10.1101/2022.12.03.518904

**Authors:** Yoojin C. Lee, Tara D. Richards, Dalia A. Fantini, David J. Kaczorowski, Bryan N. Brown, Julie A. Phillippi

## Abstract

Although *in vitro* modeling systems are becoming increasingly advanced, the complex pathophysiology of aortic diseases remains a challenge to mimic and adequately replicate. Biomechanical weakening of the vessel wall, medial degeneration and remodeling are all hallmarks of aneurysmal diseases via incompletely understood mechanisms. Understanding what factors disrupt the multi-layer biology of large blood vessels during the progression of aneurysmal disease can aid in the unmet clinical need to slow or halt disease progression. In particular, the microvascular network of the vasa vasorum provides the primary blood supply to the outer aortic wall and is a key component of inter-layer vascular health. Different origins of the vasa vasorum correspond to the anatomically specific functions of the aortic regions, which can further pertain to the differing origins of vascular wall cells and putative differences in the composition of extracellular matrix (ECM). Biologic scaffolds produced from ECM are useful biomaterials to understand biological processes and address wound healing, stem cell differentiation, and angiogenesis for both *in vitro* and *in vivo* disease models. In the present study, we investigated putative differences in composition and structure between ascending and descending aorta-derived ECM to better understand intra- and inter-layer cell-matrix interactions relevant to vasa vasorum function in the aorta. Ascending and descending aortic ECM (AECM) hydrogels were shown to retain bioactivity and influence contractility of human vasa vasorum-associated pericytes. A comprehensive understanding of the effect of layer-specific ECM on cells in different aortic regions could help uncover novel disease mechanisms.

## INTRODUCTION

As the largest artery in the body, the aorta is a unique vessel that is divided anatomically into sections: ascending aorta, aortic arch, and descending aorta, which is further defined into the descending thoracic aorta and abdominal aorta. Given the varied biological and hemodynamic environments that each division of the aorta is exposed to, aortic wall compositions are unique and varied to each anatomic region. The development of the aorta is a complex process that begins during the 3rd week of gestation and progresses during embryogenesis.^1^ Different regions of the aorta vary in origin and embryological process, where the ascending aorta originates from the primordial aortic sac and truncus arteriosus (TA), the aortic arch from the aortic sac along with a pair of right and left branches, and the descending aorta from the fusion of the right and left dorsal aortas.^2,3^ In addition, the aortic wall is composed of three distinct layers (i.e., tunica intima, media, and adventitia), which furthers the biological complexity of the aorta.^4^

The aorta can become diseased in all locations stemming from lifestyle (abdominal aortic aneurysm, atherosclerosis), congenital malformations (i.e., aortic coarctation) or genetically-triggered (e.g., Marfan, Loeys-Dietz or Ehlers-Danlos syndromes), bicuspid aortic valve-associated, familial, idiopathic, or age-related degenerative ascending thoracic aortic aneurysm. Thoracic aortic aneurysm is a life-threatening disease that is a leading cause of death in the United States.^5^ Aortic aneurysms are often silent and asymptomatic, leaving patients at an increased risk for acute dissection and/or rupture. Understanding the underlying biology of aortic wall disruptors at onset and during progression of aneurysmal disease can aid in the unmet clinical need to slow or halt the disease after diagnosis. The three layers of the aortic wall differ in composition of both extracellular matrix (ECM) and resident cells. The inner layer, the tunica intima, is the endothelium of the vessel and contains endothelial progenitor cells. The middle layer, the tunica media, is mostly composed of elastic fibers surrounding a muscular layer and smooth muscle cells (SMCs). The outermost layer, the tunica adventitia, is composed of collagen and elastin fibers, and a diverse cell population that include fibroblasts and perivascular cells. The adventitia provides substantial biomechanical support and tensile strength for vessel homeostasis. In addition, the microvascular network known as the vasa vasorum that provides the primary blood supply to the adventitia and outer aortic media is a key component in the dynamic environment of vessel homeostasis and disease.

The vasa vasorum (“vessels of the vessels”) plays an essential role in maintaining homeostasis in the aorta by delivering oxygen and nutrients to the inner layers of the tissue.^6^ In the ascending aorta, the vasa vasorum originate from the coronary and brachiocephalic arteries^7^ whereas in the descending aorta, the vasa vasorum originate from the intercostal arteries.^8^ The different origins of vasa vasorum correspond to the tissue and anatomically specific functions of the aortic regions, which can further relate to the developmental origin of cell types (e.g., endothelial cells, SMCs, pericytes) and ECM of the tissue. Cellmatrix interactions are important to understand for modeling diseases but can be challenging for complex pathophysiological systems such as the aorta.

The ECM is a complex reservoir of bioactive components and is unique in its specific aortic regions and layers, composed of varying amounts of collagen, elastin, laminin, proteoglycans, glycoproteins. Biologic scaffolds produced from ECM are useful biomaterials to understand biological processes and address wound healing, ^9,10^ stem cell differentiation, ^11^ and angiogenesis ^12,13^ and as *in vitro* and *in vivo* disease models. Therefore, to better understand cell-matrix interactions and the involvement of the vasa vasorum, in the present study we sought to identify differences among layer-specific aortic ECM (AECM) hydrogels derived from ascending and descending porcine aorta. With a better understanding of the influence of AECM on human perivascular cells, AECM hydrogels may provide a better treatment approach for aortic diseases. The development of new *in vitro* models of aortic biology will require a comprehensive understanding of the influence of layer-specific ECM on cells in different aortic regions, which is also expected to help guide the development of better medical treatments for aortic aneurysms after diagnosis.

## MATERIALS AND METHODS

### Preparation of aortic ECM (AECM)

Frozen porcine aortas were purchased commercially (Tissue Source, LLC, Lafayette, IN, USA) and thawed at 4°C overnight. The ascending (from the heart to the beginning of the aortic arch, approximately 4 cm in length) and descending (from the end of the aortic arch to 8 cm in length) portions of the aorta were divided. The adventitia (ADV) and media (MED) were mechanically delaminated and decellularized with a combination of detergents and washes with agitation (300 rpm) at room temperature. Ascending and descending AECM were prepared by decellularization similar to as described previously with modifications.^14^ The decellularization process was as follows: 0.25% trypsin for 6 hours, 3 washes in dH_2_O (15 minutes each), 70% ethanol (overnight), 3% H_2_O_2_ (15 minutes), 2 washes in dH_2_O (15 minutes each), 1% Triton X-100/0.25% EDTA/0.49% Tris (w/v) (6 hours), 1% Triton X-100/0.25% EDTA/0.49% Tris (w/v) (overnight), and 3 washes in dH_2_O (15 minutes). AECMs were sterilized in 0.1% PAA/4% ethanol with agitation (300 rpm) followed by 2 washes in 1X PBS (15 minutes) and 2 washes in dH_2_O (15 minutes). AECMs were frozen at −80°C followed by lyophilization. Lyophilized ADV ECM was incubated twice in a 2:1 chloroform/methanol solution (2 hours) and washed 2 times in dH_2_O (30 minutes) to further remove lipids. All washes were performed at room temperature. ADV ECM was re-frozen at −80°C and lyophilized a second time. Both ADV and MED ECM were milled into a fine powder using a Wiley Mill with a #40 screen.

### Validation of decellularization

Decellularization for porcine AECMs met the stringent criteria for decellularized ECM.^15^ Residual dsDNA content was quantified from 15 mg of powdered ECM using the QIAamp DNA Mini Kit (QIAgen, Germantown, MD) followed by Qubit 2.0 (ThermoFisher Scientific) according to the manufacturers’ instructions. DNA content was assessed by gel electrophoresis on a 1% agarose (ThermoFisher Scientific) gel containing 0.003% (v/v) ethidium bromide (Sigma Life Science, St. Louis, MO) and visualized under UV light with a Chemidoc XRS Bioimaging Station (Bio-Rad, Hercules, CA).

### Preparation of aortic ECM hydrogels

AECM hydrogels were formulated by blending pepsin digested ECM with Type I bovine collagen (PureCol®, Advanced Biomatrix) to the desired final concentration (1 mg/mL AECM, 2.5 mg/mL collagen). The mixture neutralized to a pH approximately 7.4 in a solution of 10X PBS, 1X NaOH, and 0.1 N NaOH. All hydrogels were prepared on ice.

### Rheology

The viscoelastic properties of pepsin digested AECM and formed hydrogels were determined by rheology using a 40-mm parallel plate rheometer (AR2000 EX, TA Instruments) as previously described and according to the American Society for Testing and Materials Standard F2900-11.^16–18^ Neutralized AECM digests were prepared on ice (“pre-gel”) and tested immediately. One milliliter of “pre-gel” was loaded onto the rheometer with a parallel plate geometry pre-cooled to 4°C. Mineral oil was used to minimize evaporation of the sample during testing. A series of rheological tests were performed in the following sequence (n=3 replicates): a steady-state flow test at 4°C to determine the viscosity profile of the “pre-gel” at a range of shear rates (0.1 to 1000 s^-1^), a rapid temperature ramp from 4°C to 37°C (1°/s) to allow gelation of the “pre-gel”; an oscillatory time sweep at 37°C to measure the maximum storage modulus *(G’),* maximum loss modulus *(G”),* and gelation kinetics by applying a small oscillatory strain (0.5% at a frequency of 1 rad/s); an oscillatory frequency sweep at 37°C to determine *G’*, *G”,* and complex viscosity *(η*(ω))* over a range of angular frequencies *(ω)* (0.1 to 100 rad/s) of the fully formed hydrogel by applying an oscillatory strain (0.5%). Data was extracted using Trios software (v4.3.0) and exported to GraphPad Prism (version 9.0) for analysis.

### Turbidimetric gelation kinetics

Turbidimetric gelation kinetics was determined for AECM hydrogels as described previously.^19^ One hundred microliters of neutralized pepsin-digested ECM [1 mg/mL ECM + 2.5 mg/mL Type I bovine collagen (PureCol®, Advanced Biomatrix)] were added in triplicate to 96-well plates. Optical density readings were obtained every 2 minutes at 405 nm for up to 2 hours using a spectrophotometer (TECAN, Germany) pre-heated to 37°C. Normalized absorbance (NA) was determined by established methods published elsewhere.^19,20^ NA was calculated at 2 minute intervals using the equation:

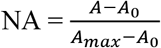

where A represents the absorbance reading at a given time point, A_max_ represents maximum absorbance, and A0 represents the initial absorbance.

### Characterization of aortic ECM hydrogel ultrastructure

Morphological ultrastructure of AECM hydrogels was assessed using scanning electron microscopy (SEM) and picrosirius red staining. Five hundred milliliters of neutralized pre-gels were placed in 1.38 cm inner diameter stainless steel annular rings and placed in a 37°C incubator for 2.5-3 hours for gelation. For picrosirius red staining, ECM hydrogels were fixed in 4% paraformaldehyde for 24 hours at room temperature, then washed in PBS and stained according to the manufacturer’s instructions (Polysciences, Warrington, PA). Samples were imaged by polarized light microscopy using a TE-2000-E2 (Nikon) and NIS Elements software (AR 5.21.02). For SEM, hydrogels were fixed in 2.5% glutaraldehyde (Electron Microscopy Sciences, Hatfield, PA) for 1 hour followed by 3 washes in 1X PBS (15 minutes each) and stored at 4°C. Hydrogels then underwent dehydration in a graded ethanol series (30%, 50%, 70%, 90%, 100% ethanol) for 15 minutes each. Samples were washed an additional 3 times in 100% ethanol for 15 minutes each and dried using a Leica EM CPD030 critical point dryer (Leica Microsystems, Buffalo Grove, IL) with CO_2_ as the transitional medium. Samples were coated with a 4.5 nm-thick gold/palladium alloy coating using a 108 Auto sputter coater (Cressington Specific Instruments, Watford, England) and imaged with a JEOL JSM6330F scanning electron microscope (JEOL, Peabody, MA).

### Elastin assay

Elastin was quantified in ascending and descending AECM using the Fastin assay kit (BioColor, UK) according to manufacturer’s instructions. Three sequential extractions of thirty milligrams of lyophilized AECMs were performed using 0.25 M oxalic acid. Each extraction was quantified separately for data analysis.

### Glycosaminoglycan content assay

Glycosaminoglycans (GAGs) were quantified in AECMs using previously described methods.^21^ Specimens (10 mg) were digested using papain (125 g/mL, Sigma-Aldrich, St. Louis, MD) in 1 mL of buffer containing 100 mM/L sodium phosphate, 10 mM/L sodium ethylenediaminetetraacetic acid (EDTA), 10 mM/L cysteine hydrochloride, 5 mM/L EDTA (all Sigma-Aldrich, St. Louis, MD) at 60°C for 16 hours. GAGs were quantified using 1,9-dimethylmethylene blue dye (pH = 3) using a spectrophotometer detecting absorbance at 525 nm. GAG concentrations were calculated from a standard curve of Chondroitin-6-Sulfate from shark cartilage (Sigma-Aldrich). Standards were ranged 0, 1.95, 3.90, 7.81, 15.63, 31.25, 62.5, 125 μg/mL.

### Isolation and culture of aortic adventitia-derived pericytes

Human ascending thoracic aorta specimens were collected during heart transplantation with informed consent and approval of the Institutional Review Board at the University of Pittsburgh (#STUDY20040179). Upon excision, tissue specimens were placed in saline on ice and immediately transported to the laboratory. Non-aneurysmal ascending aortas (aortic diameter ≤ 44 mm) were collected from 1 male donor (2 of unknown sex). Adventitia-derived pericytes were isolated from non-aneurysmal human aortic specimens and immortalized to improve culture longevity using a lentiviral vector system to deliver HPV-E6/E7 (ABM, Richmond, British Columbia, Canada) using previously described methods.^22,23^

### Contractility assay

Immortalized aortic adventitial pericytes were embedded within hydrogels (1×10^6^ cells/mL) consisting of bovine Type I collagen gels (2 mg/mL, Advanced BioMatrix, PureCol) in the presence or absence of AECM (250 μg/mL). Cell-embedded AECM hydrogels were seeded in 50 μL quadruplicates in a 12-well plate and placed in a humidified incubation chamber at 37°C with 5% CO_2_ for approximately 3 hours. Culture medium (DMEM + 2.5% FBS + 1% penicillin streptavidin) was then added, and ECM hydrogels were gently detached from the plate using a cell scraper. Cell-embedded hydrogels were imaged using bright-field microscopy (Nikon SMZ24, 1X SHR Plan Apo Objective) at t = 0 hours and after 48 hours of culture. Degree of collagen gel compaction was calculated as the inverse of the change in gel area after 48 hours of culture as measured using NIS Elements AR 4.60 software (Nikon).

### Statistical analysis

Statical analysis was performed using a one-way ANOVA with Turkey’s post-hoc test for pairwise comparisons (GraphPad Prism, version 9.0) and a p < 0.05 was considered statistically significant. Experiments were repeated a minimum of three times with at least three different cell populations and assay replicates (n = 3) per experiment. All results in the manuscript are expressed as mean ± SEM.

## Results

### Validation of decellularization of aortic ECM

Hematoxylin and eosin (H&E)-stained sections of decellularized AECN prepared from ADV and MED revealed no visible nuclei compared to an adjacent native (non-decellularized) tissue counterpart (Fig 1A). DNA gel electrophoresis qualitatively showed no large fragments of DNA (>200 bp)^24^ for both ascending and descending ADV and MED ECM (Fig 1B). Quantification of total DNA content revealed all AECM retained <130 ng DNA/mg dry weight ECM, with ADV ECM having less remnant DNA than MED ECM for both ascending and descending aorta (Fig 1C).

**Figure 1.**
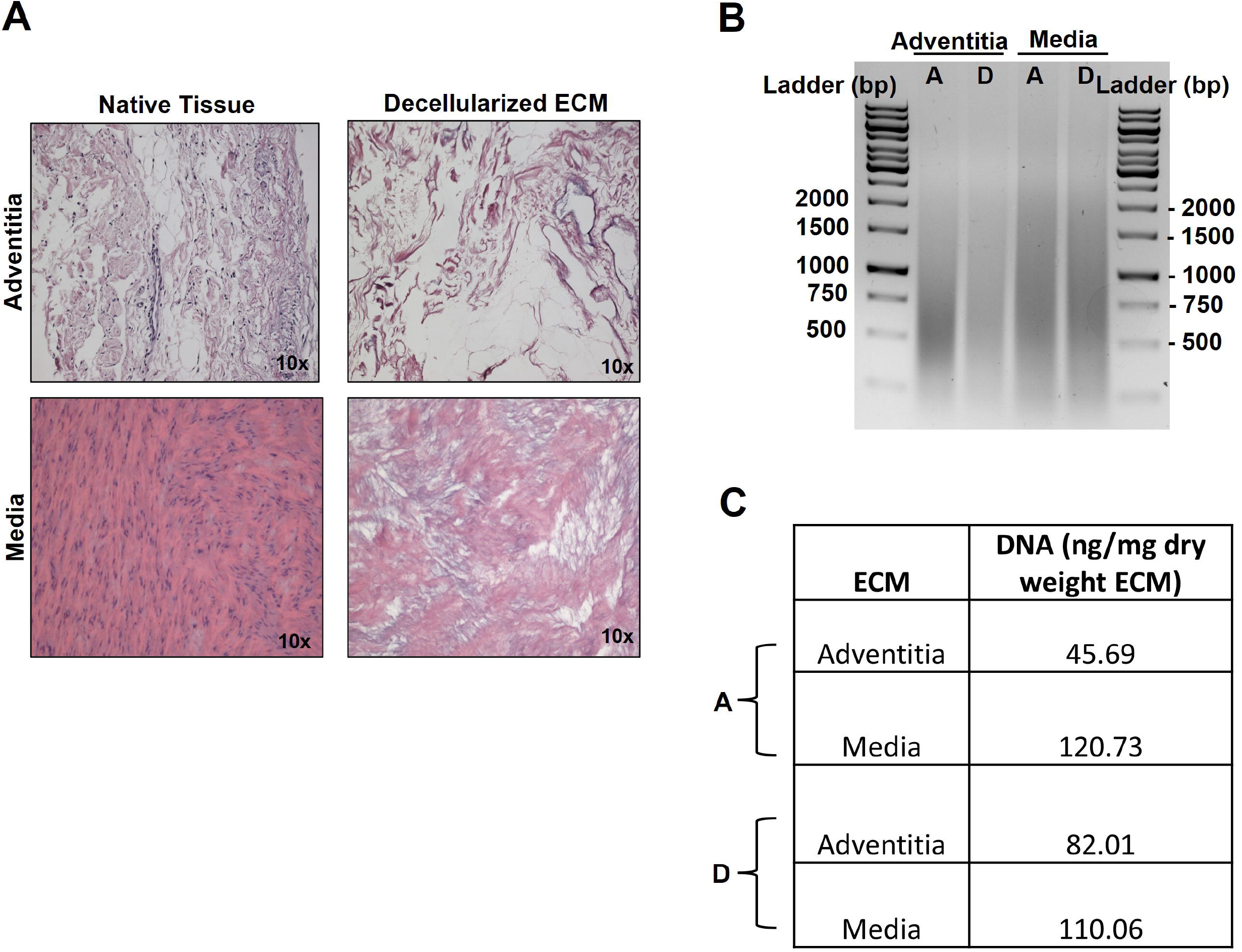
Decellularized aortic ECM. (A) Decellularization efficiency of ascending and descending aorta was validated by H&E, which showed little to no nuclei in ECM compared to native tissues. (B) DNA gel of decellularized aortic ECM did not show large DNA fragments and smears on the gel indicated small fragments of DNA (A = ascending, D = Descending). (C) Quantification of DNA determined that all aortic ECM had DNA content <130 ng/mg dry weight ECM.

### Ascending and descending aortic ECM hydrogels exhibit viscoelasticity and stability

Rheological characterization of AECM hydrogels determined that all AECM were “shear thinning,” defined as a decreasing viscosity with increasing shear rate (Fig 2A). Shear thinning of the digested ECM solution is an important property that enables injectability through a needle and syringe for future *in vivo* and clinical studies. Although there were no striking differences in viscosity between ascending and descending ADV and MED ECM hydrogels, the viscosity values for all samples were lower than 0.1 Pa*s, which indicate suitability for injection. Hydrogels formed at 37°C from ADV ECM exhibited higher maximum storage *(G’)* and loss *(G”)* modulus values when compared with MED ECM or bovine Type I collagen alone, indicating a stiffer hydrogel (Fig 2B). Gelation time increased slightly with the addition of AECM compared to collagen alone (Fig 2C). All AECM-supplemented hydrogels and collagen only hydrogels achieved peak gelation within 120 minutes. All AECM-supplemented hydrogels were “stably formed” as defined as the moduli *(G’* and *G”)* being weakly dependent on the angular frequency (*ω*), and hence the complex viscosity *(η*(ω))* was shown to be inversely related with frequency (Fig 3).^17,20^ Further, the *G’* was approximately 10-fold greater than the *G”* over the angular frequencies tested (0.1 to 100 rad/s) at low angular frequencies (≤10 rad/s) for all hydrogels.

**Figure 2.**
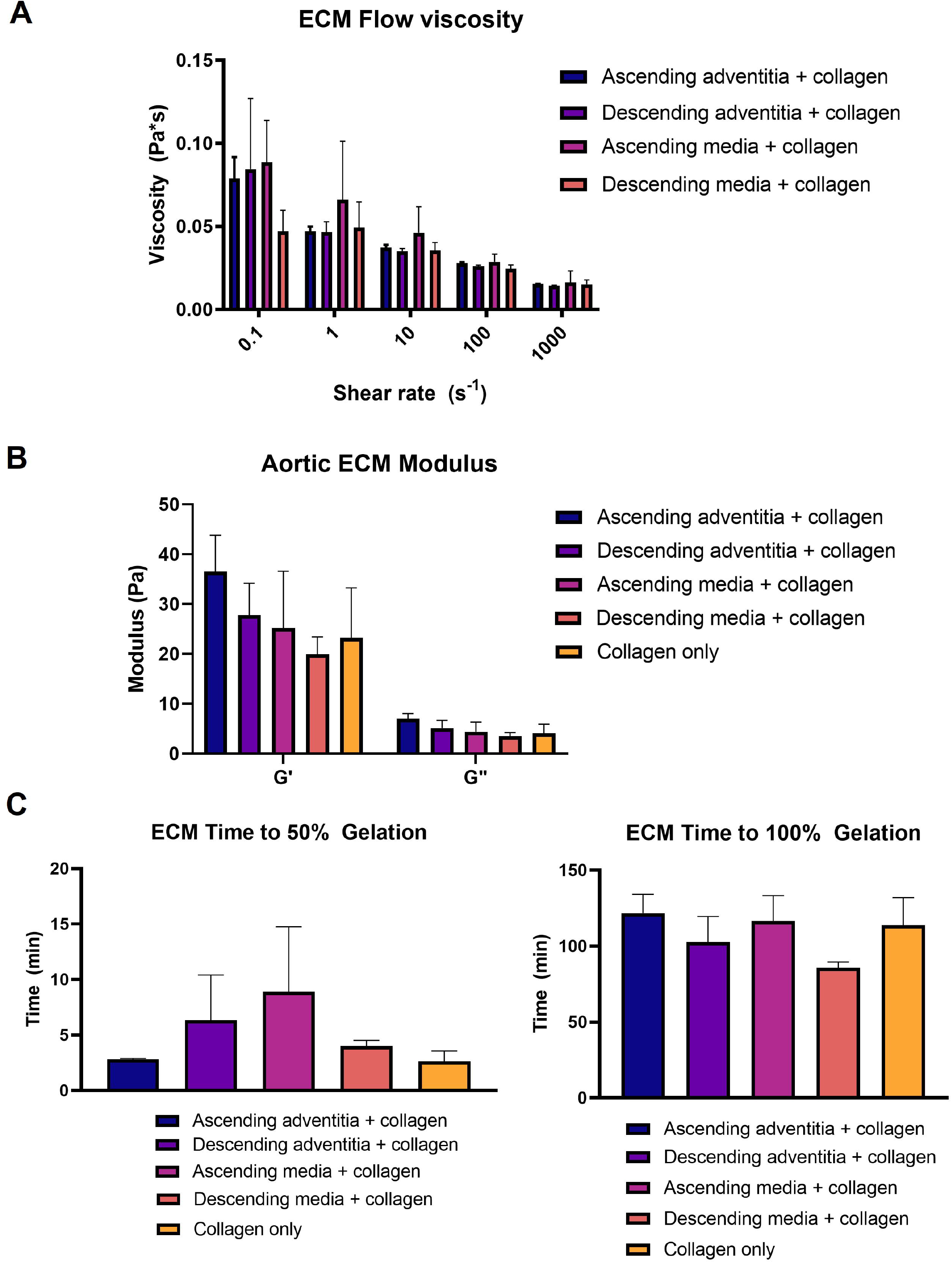
Viscoelastic properties of aortic ECM. **(A)** ECM pre-gel viscosity were “shear thinning” as defined as a decreased viscosity with increasing shear rate. **(B)** All aortic ECM hydrogels were “stably formed” as defined as the G’ storage modulus being approximately 10-fold greater than the G” loss modulus over the angular frequencies tested (0.1 to 100 rad/s) **(C)** Time to gelation (n=3).

**Figure 3.**
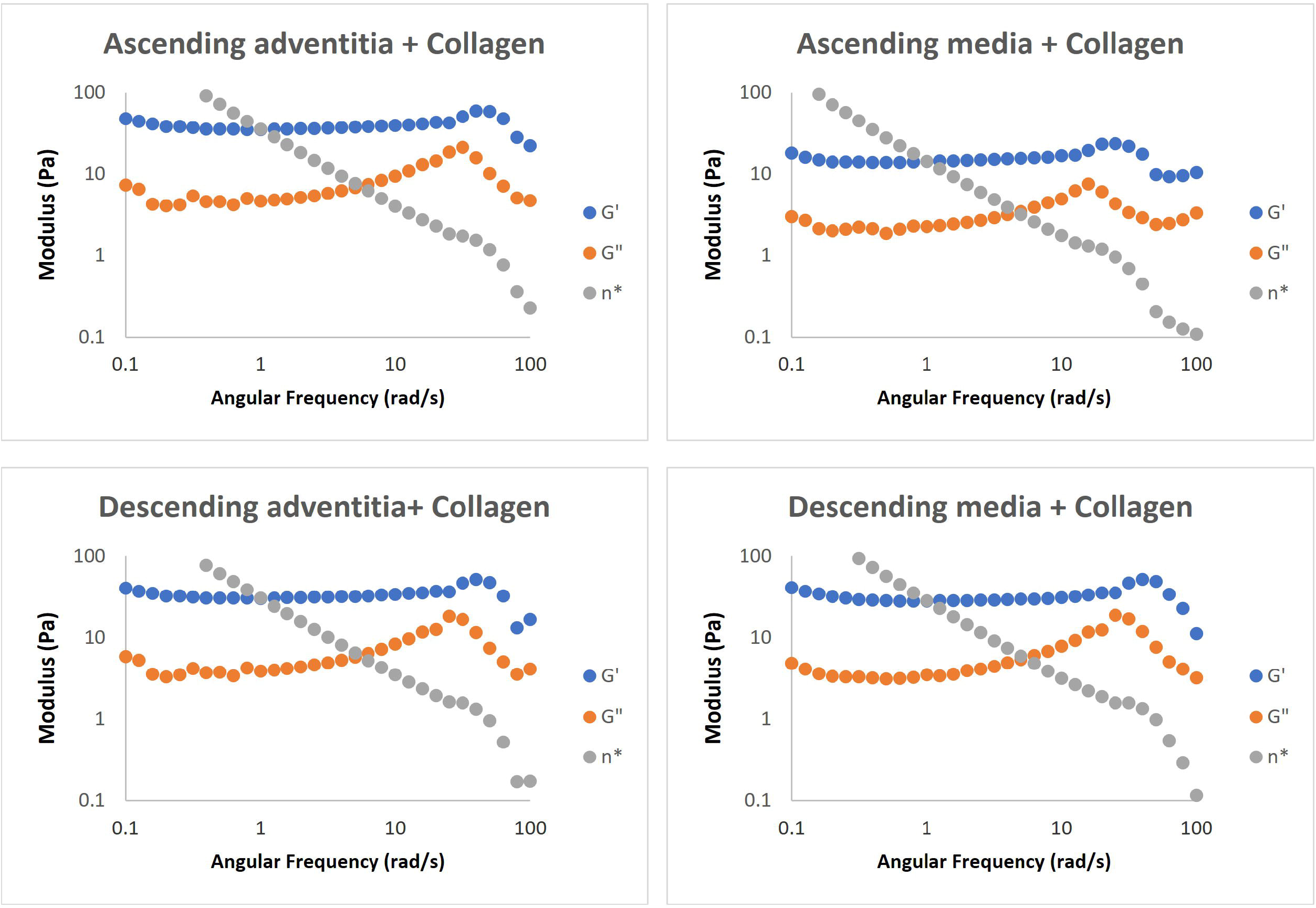
Aortic ECM hydrogels form stable hydrogels. All aortic ECM hydrogels with collagen form stable hydrogels as defined as the moduli (*G’* and *G”*) being weakly dependent on the angular frequency (*ω*), and hence the complex viscosity *(η*(ω))* is inversely related with frequency.

Optical density and normalized absorbance calculations from turbidimetric gelation kinetics revealed a logarithmic curve during the gelation period at 37°C. Peak gelation for all hydrogels were achieved within 60 minutes, and the addition of either ascending or descending AECM accelerated gelation by approximately 30 minutes (Fig 4).

**Figure 4.**
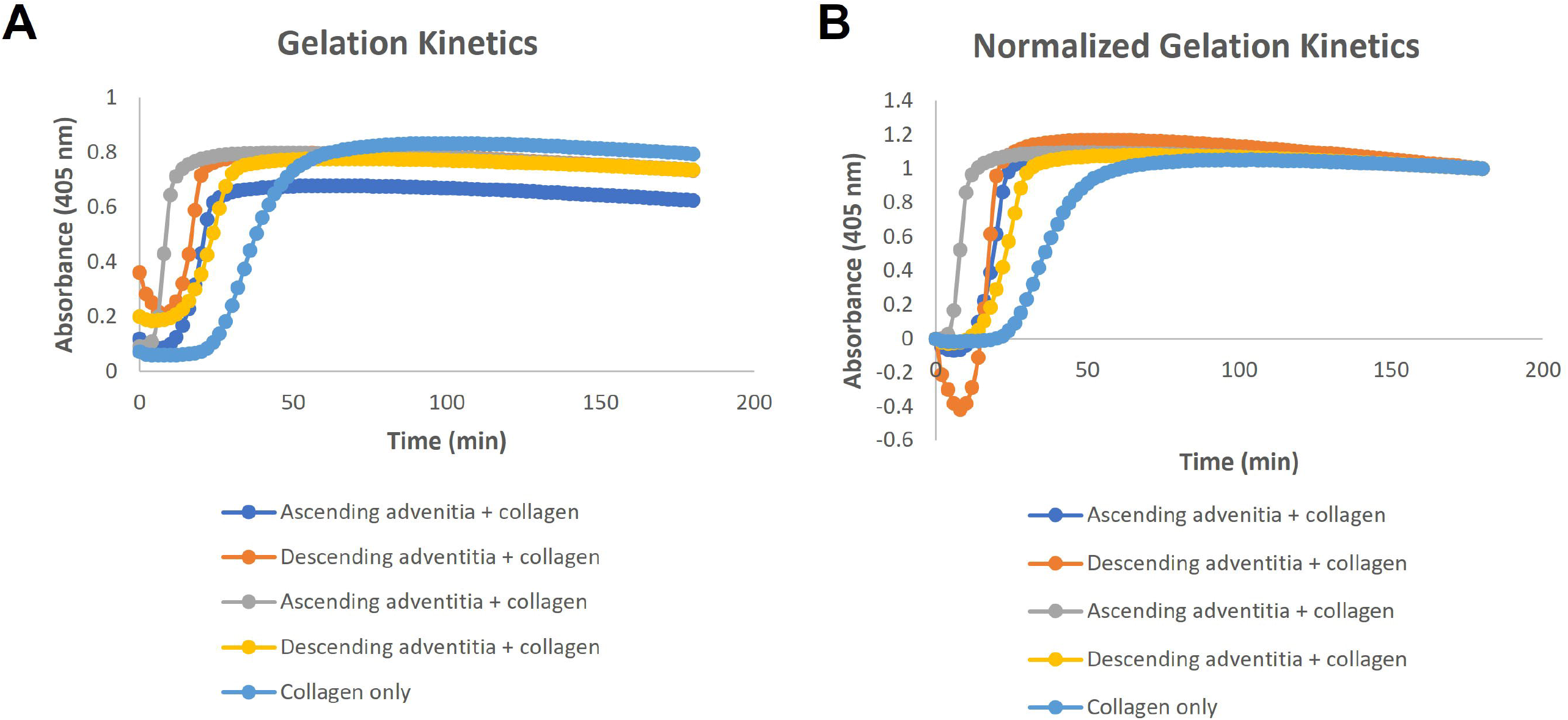
Turbidimetric gelation kinetics of aortic ECM. Optical density (A) and normalized absorbance (B) of hydrogels from ascending and descending aorta-derived ECM revealed a logarithmic curve during the gelation period at 37°C. Peak gelation of ECM hydrogels was achieved within 30 min while collagen alone gelled within 1 hour.

### Aortic ECM hydrogels exhibited layer-specific differences in ultrastructure and composition

The collagen fiber ultrastructure of aortic ADV hydrogels were similar macroscopically to collagen only hydrogels as revealed by Picrosirius red staining (Fig 5A). MED hydrogels appeared to have a more porous ultrastructure of collagen fibers. Scanning electron microscopy (SEM) of aortic MED ECM hydrogels revealed collagen fiber bundles, whereas ADV ECM exhibited predominantly single fibers in the ultrastructure (Fig 5B). The elastin content of both ascending and descending ADV ECM was lower than in MED ECM in each of the extractions with most elastin extracted in the first solubilization step (extraction 1) (Fig 6A). There was no detectable elastin in the first extraction of the media for either the ascending or descending aorta. Elastin was extracted from medial tissues only in the second and third solubilization rounds. There were no differences observed between ascending and descending ECM for either ADV (p = 0.25) or MED ECM (p = 0.19) as analyzed by paired t-test. For GAG content, ascending ECM contained less GAGs when compared with descending for both ADV and MED ECM (p < 0.0001). Overall, ADV ECM contained less GAGs than MED ECM, with significant differences between the ascending and descending regions (p < 0.0001) (Fig 6B).

**Figure 5.**
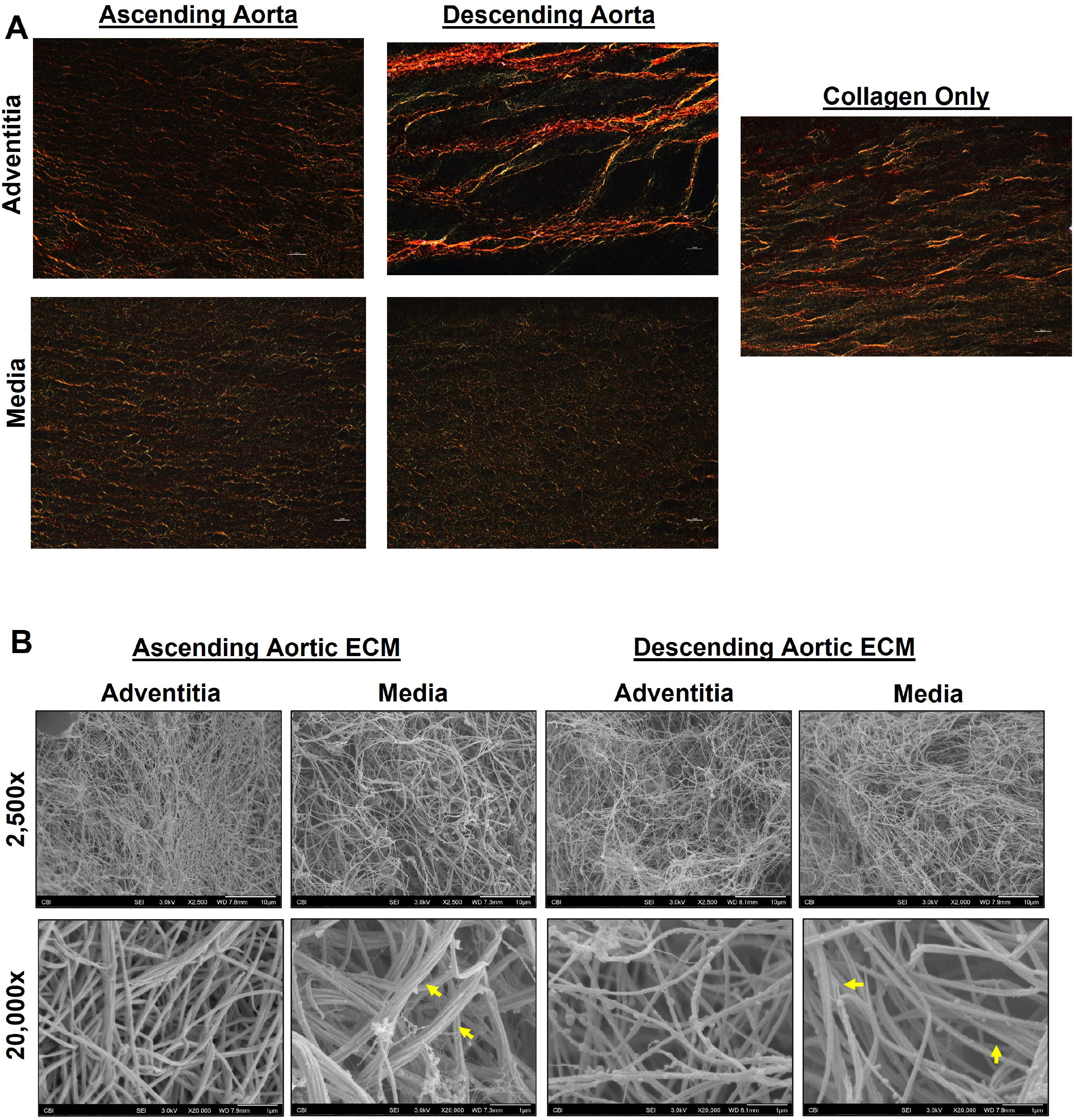
Collagen composition and ultrastructure of aortic ECM hydrogels. **(A)** Picrosirius red staining of aortic ECM hydrogels (60X magnification) **(B)** SEM of ECM hydrogels showed striking differences between the adventitia and media samples of aortic ECM. Medial aortic ECM hydrogels showed collagen fibers as more bundles (yellow arrows) than singular fibers, which were more present in adventitial ECM.

**Figure 6.**
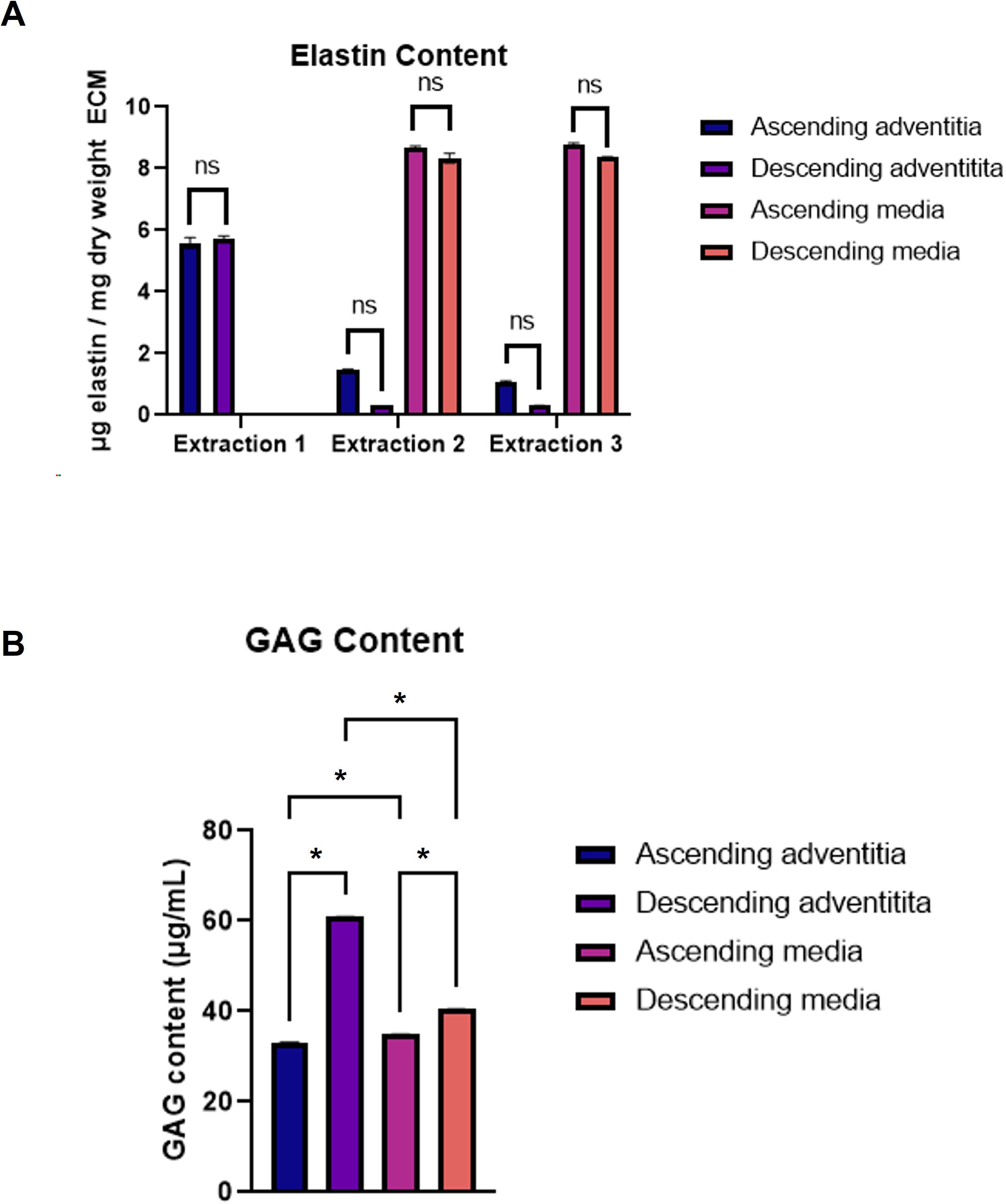
Composition of aortic ECM. **(A)** Sequential extractions of elastin from aortic ECM showed elastin content of both ascending and descending ADV ECM to be lower than in MED ECM in each extraction. No elastin was detected in the first extraction of MED ECM for either ascending or descending aorta. Elastin was extracted from MED ECM only in the second and third solubilization rounds. No differences were observed between ascending and descending ECM for either ADV or MED ECM. **(B)** Quantification of GAGs demonstrated that ascending ECM contained less GAGs compared to descending for both ADV and MED ECM. ADV ECM contained less GAGs than MED ECM, with significant differences between the ascending and descending regions. \**p*<0.0001 ns= not significant.

### Aortic ECM hydrogels are bioactive and influence aortic cellular response

To determine bioactivity of AECM, ascending and descending AECM were used as treatments in contractile assays using immortalized normal human aortic adventitial pericytes. Results were variable between the patient lines, where contractility was more robust in Patient 1 cell population than Patient 2, while Patient 3 cells failed to demonstrate any induced contractility (Fig 7).

**Figure 7.**
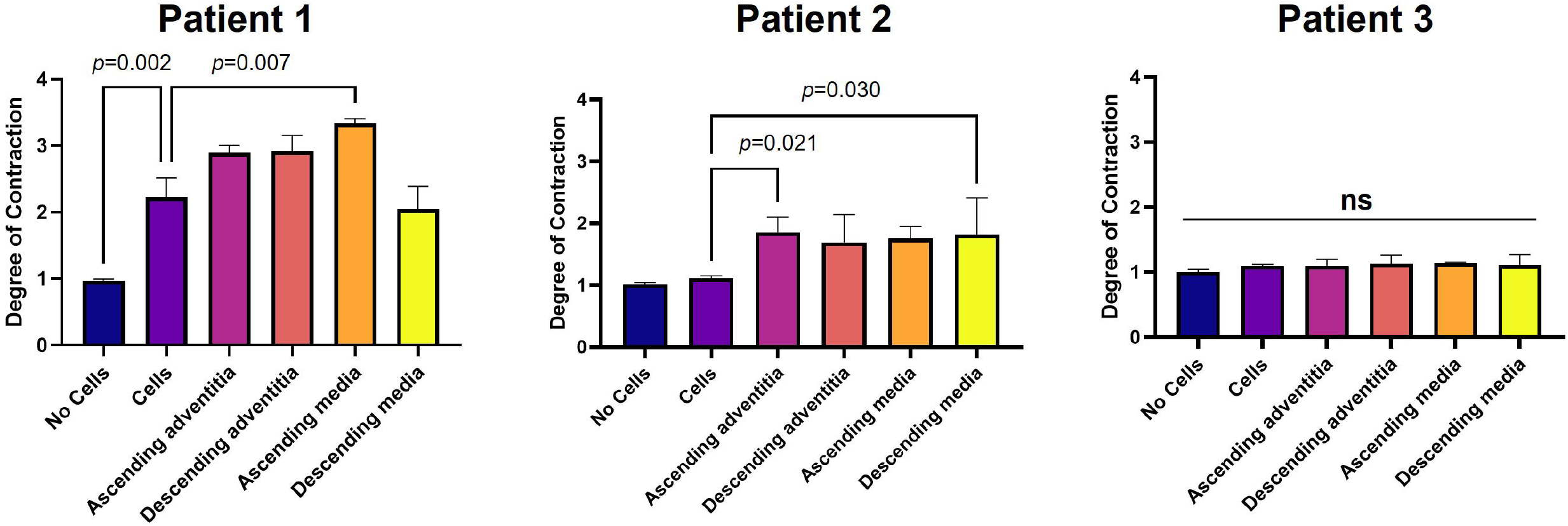
Aortic ECM variably affect vasa vasorum-associated pericyte contractility. Ascending and descending aortic ECM were used as treatments in contractile assays using immortalized (Imm) normal human vasa vasorum-associated pericytes derived from the adventitia. Results were variable, where contractility was more robust in Patient 1 cell line than Patient 2, and Patient 3 failed to demonstrate any contractility. ns= not significant.

## Discussion

The present study reports the similarities and differences in structure, composition, and bioactivity of ECM biomaterials derived from different anatomic and layers of the aortic wall. We chose to separately investigate the adventitial and medial layers of AECM from the ascending and descending porcine aorta. The aorta, the largest vessel in the human body, is composed of three distinct layers that possess specific biological functions of the tissue. The adventitia, which is the outermost, collagen-rich layer of the vessel, is composed of small blood vessels (i.e., arterial and venous vasa vasorum), lymphatics, and nerves. The tunica media is a thicker layer composed of SMCs and the ECM. The intima is the thin inner layer composed of endothelial cells and connective tissue. Among the numerous decellularization protocols published in literature, we implemented one that maintained ECM ultrastructure while removing cellular components, but also avoided harsh chemicals such as sodium dodecyl sulfate (SDS) that could potentially decrease biocompatibility and bioactivity, and destabilize the collagen network in the ECM.^25,26^ Although it has been shown to decellularize tissues successfully, SDS treatment can disrupt the triple helical domain of collagen and swell the elastin network.^27,28^ The perceived differences in the decellularization efficiency of the adventitia versus medial layer are likely due at least in part, to structural differences of the tissue, where the media is a thicker layer for the detergents to penetrate during decellularization.

There were also composition differences noted between the layers of the aorta, a vessel that must withstand the highest hemodynamic forces in the body. Elastin content in the prepared AECM biomaterials reflects both layer specific and regional heterogeneity of the aorta, whereby the adventitia contains less elastin than the media, and there is less elastin in the descending thoracic aorta and abdominal aorta than in the ascending thoracic due to hemodynamic requirements.^29^ The adventitia contains less elastin when compared with the media, which can have 40 or more elastic lamellae in healthy human adults with age and aortic disease affecting the number of elastin lamellae and overall content.^30^ Our DNA gel analysis and histomorphologic assessments confirmed small fragments of remnant DNA and no visible nuclei, respectively. Despite differences in remnant DNA content for adventitia and media, we do not anticipate any deleterious effects in our *in vitro* or future *in vivo* experiments since values for DNA are similar to previously published amounts for porcine adventitia.^19^

The viscoelastic properties of AECM hydrogels are important when considering future applications for these hydrogels. It has been shown that the concentration of ECM in a hydrogel influences the viscoelastic properties of the gel.^17^ The decellularization process is important to remove the cellular components in the ECM and retain the bioactive matrix components, all the while taking care to avoid any remnant detergents, which can disrupt the gelling properties of the hydrogel.^31^ In our study, we showed that the adventitia produced a hydrogel with higher storage modulus (resulting in a “stiffer” hydrogel), which likely is because it is the insulating outer layer and provides the majority of tensile strength of the artery. The decellularization process of adventitia and medial layers of the aorta were identical, except the adventitia received an additional chloroform-methanol step to further deplete lipids. Without this additional step to remove lipids, milling adventitial ECM into a powder in its lyophilized state was suboptimal and resulted in low yield. Our decellularization protocol relies on a combination of trypsin to enzymatically remove cells, as well as alcohol (75% EtOH), a non-ionic detergent (Triton X-100), acid (PAA), and chelator (EDTA) in sufficient amounts of exposure time to minimize disruption to the ECM. These agents have consistently achieved effective decellularization in numerous tissues, including the aorta.^32–34^ Levy *et al,* showed ethanol treatment of tissue cross-links collagen thus influencing the fiber ultrastructure.^35^ It has been shown that GAGs and ECM constituents interact and play a role in ECM recruitment the biomechanics in the arterial wall.^36^ Although the decellularization process depletes GAGs and other ECM-related molecules, much of the sulfated GAGs have been shown to be retained to support binding of growth factors such as basic fibroblast growth factor (bFGF).^37^ We also avoided using SDS in our protocol since it has been shown to markedly reduce GAGs and presence of growth factors in the ECM.^26,38^ Interestingly, decellularized adventitia of the descending thoracic aorta exhibited the higher GAG content when compared with the media of both the adventitia and media of the ascending and the media of the descending thoracic aorta.

A substantial advantage of using a hydrogel is that it can be tailored to contain or omit various factors (e.g., ECM proteins, GAGs, and growth factors) by the type of source tissue. The main difference in the ultrastructure of AECM hydrogels was layer-specific with noted distinctions in the adventitia, and media. SEM revealed that in adventitia, the network of collagen appeared to be more singular fibers whereas in media, the collagen fibers were noted mostly in bundles. Since collagen bundles were observed for both the ascending and descending aortic media, this is likely due to the compositional differences between the adventitial and medial layers of the aorta. The adventitia is the outer superficial layer of the vessel and is composed of collagen and elastic fibers. The media, which is the densest layer of the aorta, is composed of elastic lamellae as well as collagen, which maintain the structural homeostasis of the vessel. It has been shown that SMCs anchor to the lamellar layers of elastin to form the aortic medial ultrastructure known as “contractile-elastic units.”^39^

AECM hydrogels are bioactive and vary in influencing contractile activity in human adventitial pericytes. The variable response of vasa vasorum-derived pericytes to AECMs may be due to patient-to-patient variability or potential heterogeneity of the population.

In conclusion, this work highlights the importance of tailoring decellularization strategies for different layers of the blood vessels wall. Intriguingly, ECM hydrogels recapitulate distinct differences in composition and ultrastructure that are unique to each layer. Importantly, ascending and descending AECM hydrogels retain bioactivity to influence human perivascular cells and can potentially be used as disease models for investigating arterial diseases such as aortic aneurysm. A comprehensive understanding of the influence of layer-specific ECM on cells in different aortic regions could help uncover novel disease mechanisms and serve as less invasive treatments for aortic aneurysms.

## Acknowledgements and Funding

This work was supported by NHLBI under award #HL131632 (J.A.P.), the Commonwealth of Pennsylvania, University of Pittsburgh Physicians (J.A.P.), the Department of Cardiothoracic Surgery (J.A.P), and the UPMC Pellegrini Chair in Cardiothoracic Surgery (J.A.P.). The authors thank Katelyn Osborn and Sarah Bill for their assistance with patient enrollment.

## Notes

### Competing Interest Statement

The authors have declared no competing interest.

